# Temporal Insights into Molecular and Cellular Responses during rAAV Production in HEK293T Cells

**DOI:** 10.1101/2023.11.02.565245

**Authors:** Alok Tanala Patra, Evan Tan, Yee Jiun Kok, Say Kong Ng, Xuezhi Bi

## Abstract

The gene therapy field is actively pursuing cost-effective, large-scale production of recombinant adeno-associated virus (rAAV) vectors for therapeutic applications, which demand high dosages. Enhanced yield is essential but presents technical and cost challenges. Strategies like suspension cell culture, transfection optimization, and cultivation conditions have shown moderate success but fall short for large-scale applications. In this study, we explore the host cell proteome of HEK293T cell lines used for AAV2 production under different transfection conditions (standard, sub-optimal, optimal) using SWATH-MS. To understand molecular and cellular mechanisms, we created a tailored spectral library for HEK293T-AAV interactions. Our gene ontology and pathway analysis revealed significant protein expression variations, particularly in processes related to cellular homeostasis, metabolic regulation, vesicular transport, ribosomal biogenesis, and cellular proliferation under optimal transfection conditions. These conditions increased rAAV titre by 50% compared to standard protocols. Furthermore, we identified alterations in host cell proteins crucial for AAV mRNA stability and gene translation, particularly regarding AAV capsid transcripts in optimal transfection conditions. This study provides insights into cellular mechanisms during rAAV production in HEK293T cells and offers potential advancements in scalability and cost-efficiency for gene therapy vector production.

**Significance:** Generating AAV is challenging and the amount produced is limited, despite the success of gene therapy. Thus, it is necessary to enhance the productivity of host cells through various engineering techniques. The replication and transduction of viruses cause numerous modifications to the host cell’s proteome. Viruses exploit the processes of the host cell, influencing the availability, post-translational adjustments, associations, or placement of its proteins during the replication/growth cycle. Comprehending the fluctuations in biological processes during AAV replication is essential to grasping how virus-host relations shape the result of infection and viral protein expression.

## Introduction

Gene therapy has emerged as a promising treatment approach for a number of genetic diseases, certain types of cancer, and other infectious diseases. Although several types of gene delivery “vectors” such as viral or non-viral vectors are available, the recombinant adeno-associated virus (rAAV) is currently the main viral vector of choice for gene delivery to treat a variety of human diseases ^1–7^.

Adeno-associated viruses (rAAVs) are a small, non-pathogenic virus that can be used to deliver gene therapy to target tissues. They have been used in a variety of applications, including gene therapy, gene editing, and vaccine development. AAVs have been used to treat genetic diseases such as hemophilia, cystic fibrosis, and muscular dystrophy, as well as cancer. In order to ensure the successful application of AAVs, the use of recombinant Adeno Associated Virus (rAAV) as a gene therapy vector has become increasingly popular due to its ability to efficiently deliver genes into target tissues with minimal toxicity and long-term transgene expression ^8–13^. One pivotal challenge within this landscape centers on identifying optimal transfection conditions capable of augmenting the production of high-titer rAAVs containing the desired gene of interest (GOI)^14–16^. While AAV yields have demonstrated promise, the specific hurdle to surmount revolves around ensuring a substantial quantity of fully loaded AAV particles carrying the GOI. This aspect is critical for realizing the full potential of rAAVs and broadening their applications in the realm of gene therapy and related fields.

We use Design of Experiment (DoE) as a method to optimize rAAV production. DoE involves identifying key variables that affect the production process and systematically varying those variables in order to maximize yield. The goal of optimizing rAAV production using DOE is to identify the most effective combination of parameters for producing high- quality, high-yield virus particles. This includes factors such as cell line, vector design, transfection protocol, and purification conditions^17–20^. By increasing our understanding of how these various elements interact with each other during rAAV production we can develop more efficient manufacturing processes and ultimately produce higher yields from fewer resources. Additionally, DOE allows us to predict which combinations are likely to result in successful out-comes prior to conducting experiments or investing resources into full-scale manufacturing efforts - thus saving valuable time and resources in the long run.

Notably, the existing body of research has inadequately addressed the molecular and cellular dynamics of HEK293T cells during AAV production, despite their widespread use in the biomanufacturing industry. Previous studies utilizing DoE have primarily focused on AAV yield in various transient transfection conditions^21–23^, offering limited insights into the molecular dynamics underlying the conditions associated with higher AAV production.

Furthermore, some studies have concentrated solely on comparing HEK293 cell lines in non- transfection conditions versus standard transfection conditions during specific harvesting time points^24^. These investigations, however, often neglect to explore the molecular dynamics of the host cell across different time points, a crucial element for comprehending the temporal dynamics across various cellular states during the AAV production process.

Recognizing this substantial knowledge gap, our study leverages DoE to identify optimal transfection conditions based on AAV titer. In addition, we utilize the SWATH-DIA method to gain a comprehensive understanding of the host cell’s cellular states under various transfection conditions compared to the standard condition. This combined approach addresses the dearth of information regarding the molecular and cellular dynamics during rAAV production, offering insights crucial for optimizing bioprocessing strategies in the future.

In our study, we have optimized triple transient transfection conditions for the AAV2 serotype expressing GFP and formalized transfectant conditions to achieve ∼50 % higher yield than the standard transfectant that is currently available. Further, to understand the dynamic role of HEK293T cells at different cellular states post-transfection we have performed a comprehensive comparative proteome profiling of the host cell system post-transfection at 24, 48, and 72 hpt. For this study, we developed an in-house label-free HEK293T-AAV spectral library which comprised ∼7,183 protein groups. We identified about 4800 protein groups using our spectral library for a comparative proteome study. A proteome profiling and gene ontology analysis was performed for 3 different transfection conditions (including a standard plasmid molar ratio of 1:1:1) to elucidate the molecular and cellular processes of the host cell (HEK293T transfectants) that contribute towards higher AAV yield.

This study provides insights into how to improve the manufacturing of Adeno Associated Viruses (AAV) and could serve as a benchmark for other similar viruses.

## Results

To probe the impact of diverse rAAV production conditions on the proteome of the production cells, we harnessed various rAAV transfection conditions achieved through the DoE methodology. By meticulously optimizing DoE, we derived two distinct rAAV production conditions, each associated with graduated improvements in rAAV titers. Our research encompassed three specific rAAV production conditions, namely Standard, Sub-Optimized, and Optimized (as depicted in Fig. 1). These conditions afforded us the opportunity to scrutinize the proteomic landscape of HEK293T cells under various rAAV production scenarios. This comprehensive investigation is poised to yield critical insights into the intricate cellular processes pivotal for enhancing rAAV production, thus unveiling potential cellular targets for future advancements in cell engineering.

**Fig. 1.**
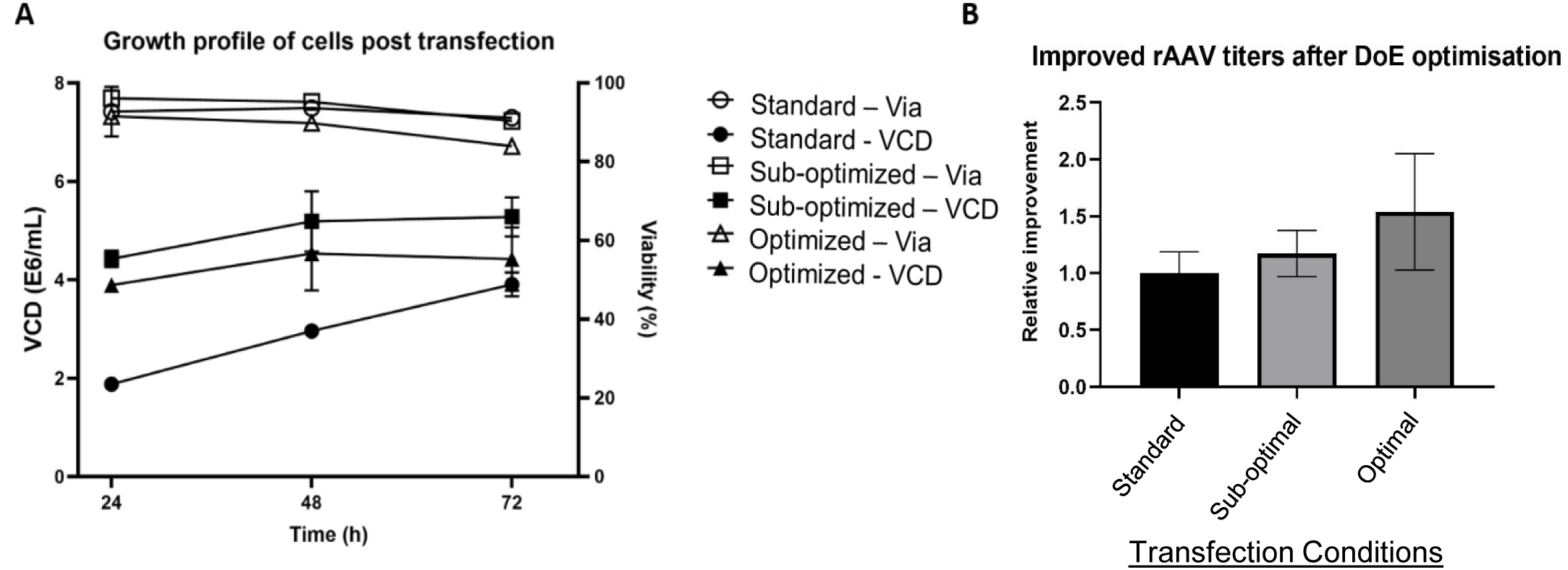
Growth profiles of cells post transfection and rAAV titers. Two rounds of DoE optimization were used to optimise the transfection process. Factors optimised included cell density, DNA concentration, plasmid ratios, and PEIMax concentration. A. Viability and Viable cell density (VCD) of cells post transfection. Cells were harvested 72 hours post transfection for rAAV titering. B. Optimisation of transfection conditions gave an ∼50% increase in rAAV titers obtained.

**Fig. 2.**
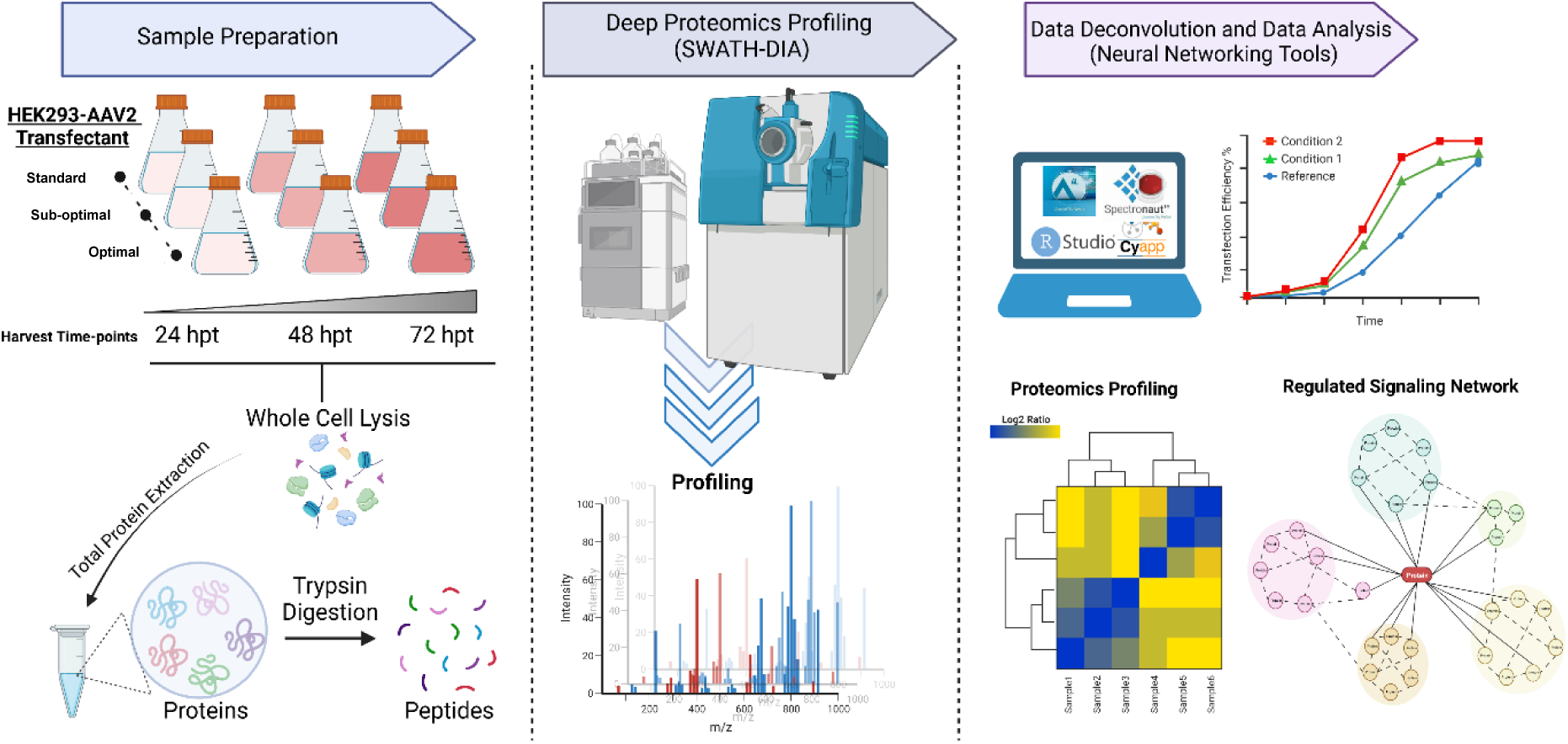
Comprehensive Spectral Library for the Analysis of the HEK293T Proteome by SWATH-MS. (A) Construction and Validation of the SWATH HEK293T Global Spectral Library. HEK293T cell samples underwent differential ultracentrifugation to isolate nuclear (NP), mitochondrial (MP), and membrane (MP) proteins. The lysates from whole cells (WCL) and subcellular fractions were digested, fractionated, and analyzed by DDA-MS. The DDA data facilitated the creation of a spectral library using Spectronaut software. The library’s utility and robustness were assessed with various HEK293T-AAV samples, enhancing sensitivity and proteome depth (B). The pie chart (C) illustrates the distribution of cellular components and biological processes within the HEK293T spectral library based on PANTHER GO-Slim terms.

### Development of a Label-Free Quantification Spectral Library Utilizing HEK293T Cell Line for AAV2 Production

The creation of the global spectral library for HEK293T cells involved the acquisition of MS/MS spectra from both unfractionated and fractionated protein samples, obtained from HEK293T cells under various conditions, including AAV2 transfections. Additionally, data from three distinct subcellular organelles were integrated: nuclear proteins (10 fractions), mitochondrial proteins (10 fractions), and membrane proteins (10 fractions). Furthermore, an extensively fractionated whole-cell protein lysate from HEK293T was included, consisting of 10 fractions. The number of proteins in the HEK293T global spectral library (with a 20% coefficient of variation cut-off and a 1% peptide false discovery rate) reached a saturation point as more DDA files were introduced during the library’s generation (as depicted in Fig. 3), suggesting that the library effectively captured the majority of proteins in the sample.

**Fig. 3.**
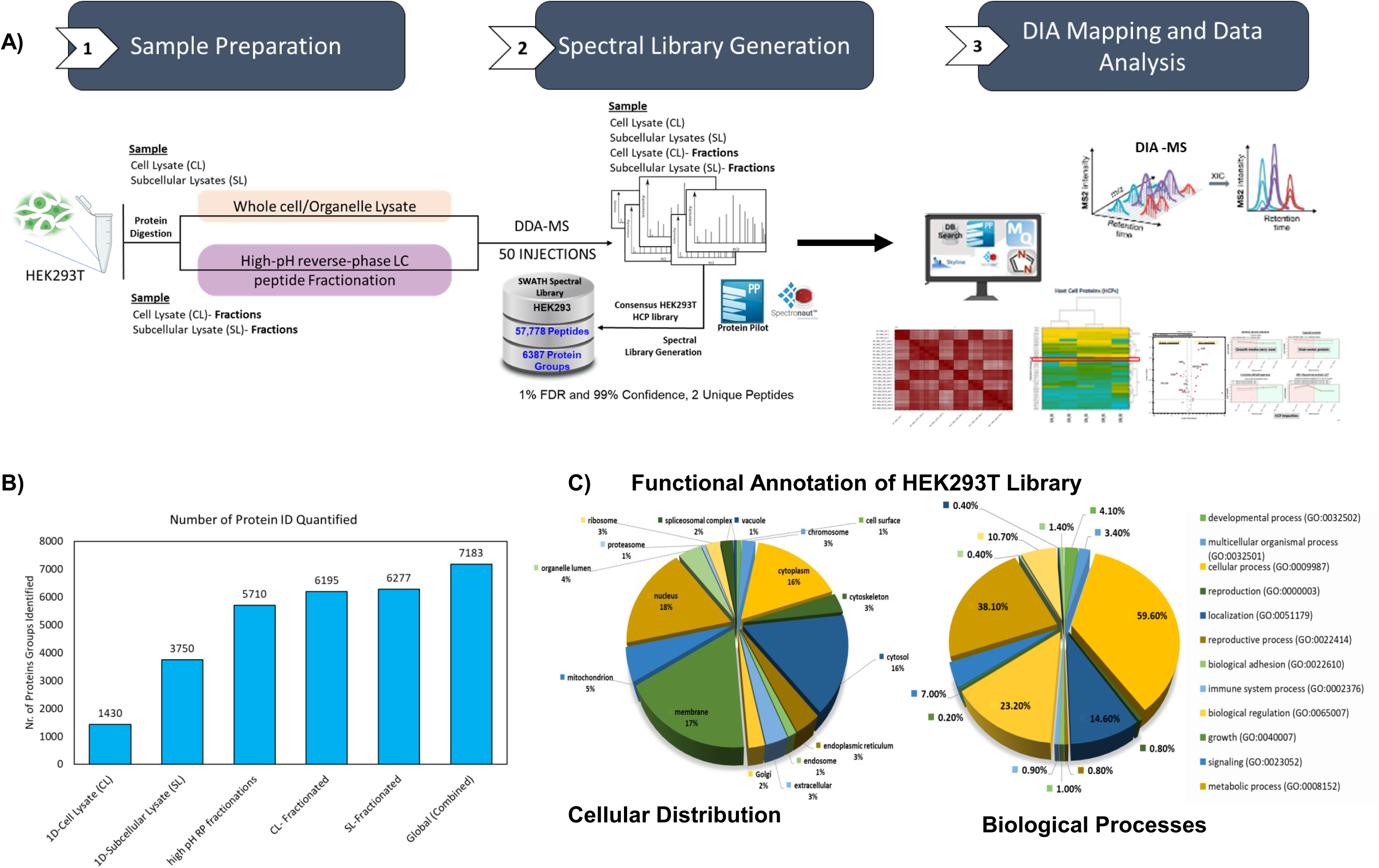
SWATH-MS Proteomics and Bioinformatics Analysis: The Experimental Workflow. **A)** The following workflow outlines the construction and use of the SWATH HEK293 global spectral library. Using our in- house protocol HEK293-derived samples were subjected to multiple rounds of processing and separations. The HEK293T cells were lysed and fractionated using differential ultracentrifugation to isolate nuclear proteins (NP), mitochondrial protein (MP), and membrane proteins (MP). The protein lysates from the whole cell (WCL) and subcellular-organelle compartments were tryptic digested, subsequently fractionated using basic reverse-phase liquid chromatography separation and subjected to DDA-MS analysis. The data from the DDA search was used to construct a spectral library in Spectronaut software. The applicability and robustness of the HEK293 global spectral library were evaluated with SWATH-MS data sets of different HEK293T-AAV samples, including WCL of different cell line transfectant samples, and using various LC-MS instrumental setups. **B)** Bar graph showing increased sensitivity and dynamic proteome depth to construct a comprehensive Spectral library for HEK293T cell line. Human Database Search for Control cell-line HEK293T (DDA runs) using Protein Pilot. The result filtration parameters used were as follows; Peptide -10lgP ≥ 15, Protein Group FDR 1.0%, and Proteins unique peptides ≥ 2. C) The pie chart showing the distribution of the cellular components and biological processes of in-house generated HEK293T spectral library. The distribution of GO terms was categorized based on PANTHER GO- Slim.

In contrast to the CL 1D library, which contained 1,430 protein groups, the 1D library constructed using cell and subcellular lysates collectively quantified approximately 3,750 protein groups, constituting a notable increase of 54.61% more proteins. By combining the CL fractions, 1D-library, and high-pH RP-separated fractions, we successfully quantified approximately 7,183 protein groups (as illustrated in Fig. 3B), surpassing previous reports ^25^. This underscores the library’s efficacy in capturing a diverse array of proteins. It’s important to note that Spectronaut 17, known for its speed and automation, was employed to create our spectral library, benefiting from features like iRT determination, optimal fragment selection from replicates, and enhanced enrichment capabilities.

Moreover, we conducted an in-depth characterization of our HEK293T SWATH spectral library. Functional annotation of the library’s proteins was performed using the PANTHER database, revealing a significant portion of these proteins associated with fundamental biological processes, particularly encompassing cellular processes, biological regulation, and metabolic processes. Regarding their cellular localization, the majority of these proteins were found within the cellular component and organelle part (as depicted in Fig. 3C).

### Profiling the Whole-Cell Proteome of HEK293T-AAV Transfectants

In this investigation, we utilized the SWATH-DIA-based label-free quantification methodology to conduct a comparative analysis of the proteome profiles among HEK293T- AAV transfectants exposed to three distinct conditions: The Standard condition, Sub-optimal condition, and Optimal condition. To enhance our analytical capabilities, we systematically constructed a curated Spectral Library for label-free quantification, utilizing the HEK293T cell line engineered to produce AAV2. This comprehensive proteome profiling was performed at multiple time points, specifically at 24, 48, and 72 hours post-transfection (hpt), across all transfectants. At 24 hpt, across all transfectants, we consistently identified approximately 4800 protein groups with an FDR of less than 0.01 with at least 1 peptide per protein (Supplementary dataset 1). Whereas for the later time points 48 and 72 hpt we noticed quite a variability in the number of protein groups identified and quantified for all the transfectant conditions (as shown in Fig. 4B). The proteomes from each biological replicate were analyzed using Two-dimensional principal component analysis (PCA), which revealed that the proteomes clustered according to the infection time point. To evaluate changes in protein abundance over the course of the infection, a differential expression analysis was conducted (using defined fold change and significance thresholds) (Fig. 4C). We used PCA (Principal Component Analysis) as a statistical tool to analyze this high-dimensional data and identify patterns or similarities between the transfectants. Here the PCA plot shows the clustering of the sample time points based on their protein profiles. Two-dimensional principal component analysis (PCA) was performed on the proteomes of each biological replicate, and it was observed that the proteomes clustered based on the infection time point. To assess variations in protein abundance during the infection, a differential expression analysis was executed, with defined fold change and significance thresholds (as shown in Fig. 3C).

**Fig. 4.**
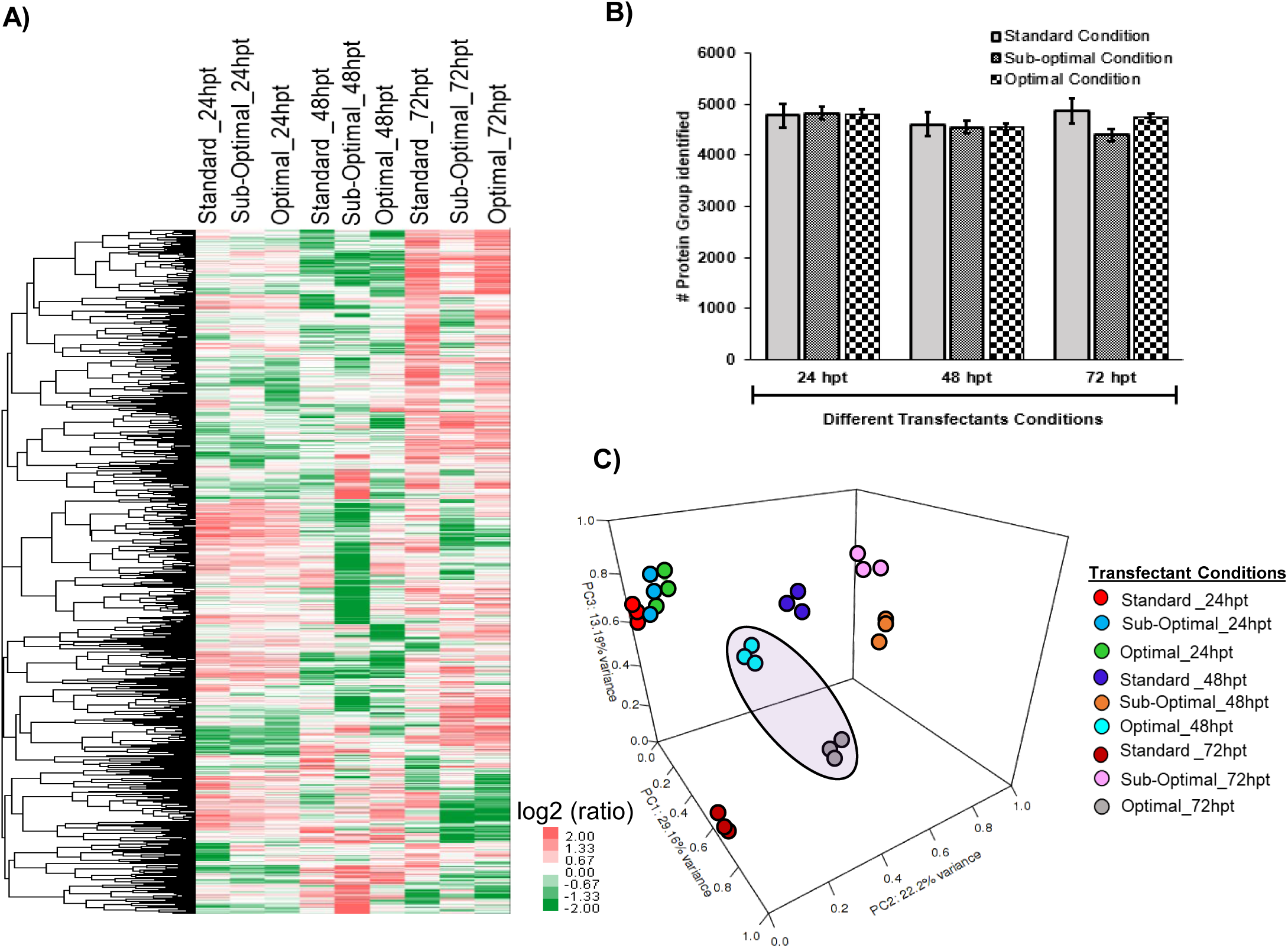
**Comparative proteomic analysis of HEK293T transfectants during AAV2 production at different time-points**. A) Overall proteomics profile for HEK293T-AAV transfectants at different time points (24, 48, and 72 hpt; hours-post transfection). The protein profile displays those protein groups which have passed a filter that removes all genes that have missing values. The relative protein abundance is represented as a heat map, with the representative proteins of each group clustered together if they exhibit a similar expression trend across the samples. The hierarchical clustering is generated using the neighbor-joining algorithm, with a Euclidean distance similarity measurement of the log2 ratios of the abundance of each sample relative to the average abundance. B) Total number of protein groups detected in HEK293T-AAV transfectant conditions (Standard, Sub-optimal, and Optimal transfectant conditions) at different time points post-transfection. C) Principal component analysis (PCA) of the individual proteomics datasets shown for all three transfectant conditions in HEK293T cell line.

### Proteomic Analysis of HEK293T Producer Cell Line under Optimal Transfection Conditions

Our central focus revolves around the elucidation of the molecular and functional transformations associated with the Optimal transfection condition, which achieved a 50% increase in titer compared to the standard transfection condition (as depicted in Fig. 1). This specific condition exhibited the highest rAAV titer at the 72- (hpt) time point in contrast to other transfection conditions. Our extensive proteome analysis, executed under this specific transfection condition, enabled the identification and quantification of approximately 5253 proteins, with a false discovery rate (FDR) of less than 0.01 (see Fig. 4B).

Moreover, at each time point (24, 48, and 72 hpt), we identified sets of differentially expressed proteins, characterized by fold changes of at least 1.5 and p-values less than or equal to 0.05. In comparison to the standard transfection condition, we observed a decrease in the abundance of 103 proteins at 24 hpt, 112 proteins at 48 hpt, and 421 proteins at 72 hpt, corresponding to the progression of rAAV production. Simultaneously, an increase in the abundance of 74 proteins at 24 hpt, 147 proteins at 48 hpt, and 139 proteins at 72 hpt was noted (as displayed in Fig. 5-7 (A)).

**Fig. 5.**
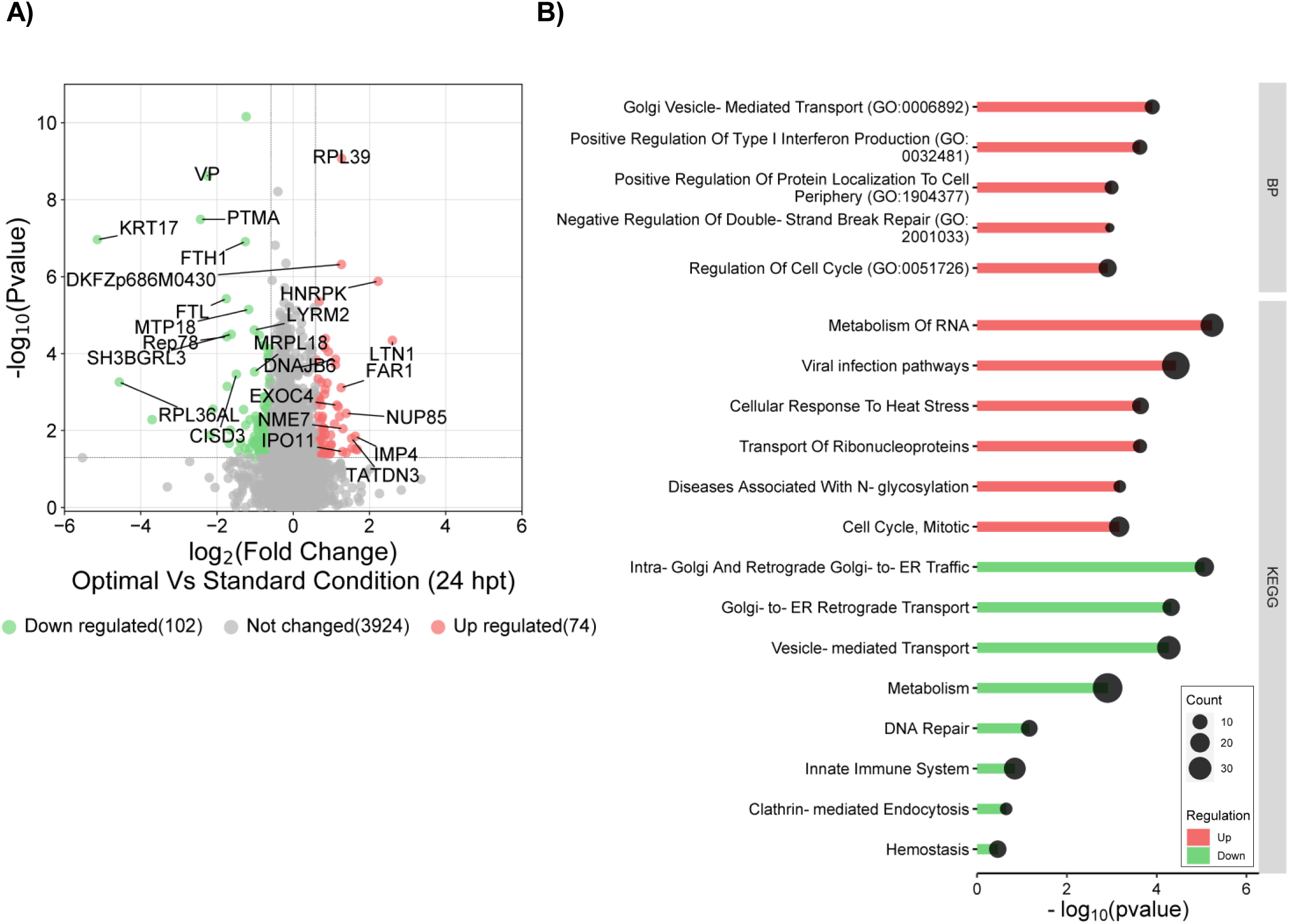
Differentially Abundant Proteins in Optimal Transfection Conditions: A 24- Hour Post-Transfection Analysis. Based on average log2 fold change and Pvalue <=0.05. Red indicated upregulation while green shows down regulation of the corresponding proteins. B) Significantly enriched Gene Ontology Biological Process and KEGG pathways revealed by Enrichr analysis of the protein expression up- (in red shade in the figure) or down- (in green shade in the figure) regulated in Optimal Transfectant condition compared Standard Transfectant condition at 24 hpt. In vertical axis is the terms, in horizontal axis is the transformed FDR (−log10PValue).

**Fig. 6.**
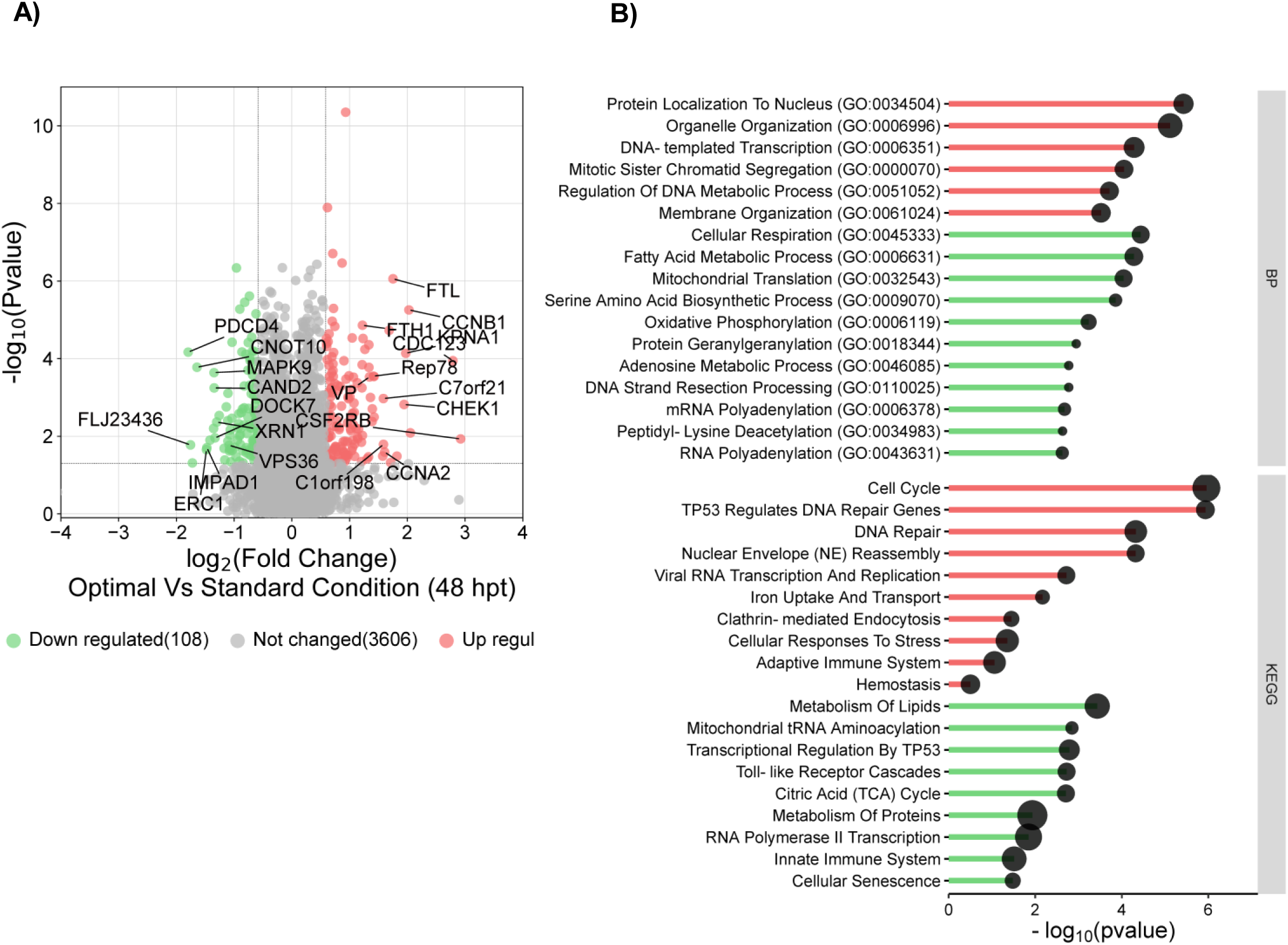
Differentially Abundant Proteins in Optimal Transfection Conditions: A 48- Hour Post-Transfection Analysis. Based on average log2 fold change and Pvalue <=0.05. Red indicated upregulation while green shows down regulation of the corresponding proteins. B) Significantly enriched Gene Ontology Biological Process and KEGG pathways revealed by Enrichr analysis of the protein expression up- (in red shade in the figure) or down- (in green shade in the figure) regulated in Optimal Transfectant condition compared Standard Transfectant condition at 48 hpt. In vertical axis is the terms, in horizontal axis is the transformed FDR (−log10PValue)

**Fig. 7.**
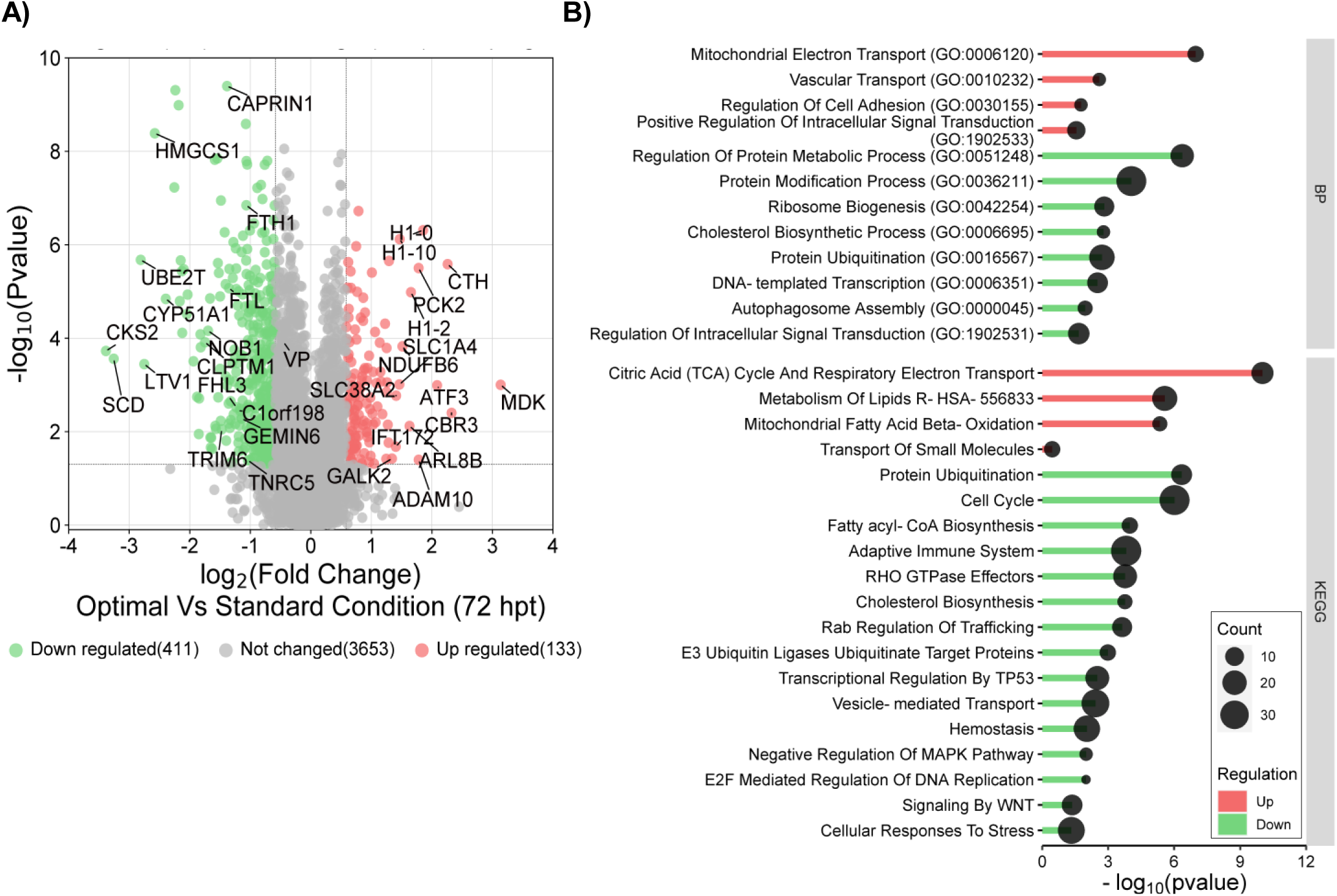
Differentially Abundant Proteins in Optimal Transfection Conditions: A 72- Hour Post-Transfection Analysis. Based on average log2 fold change and Pvalue <=0.05. Red indicated upregulation while green shows down regulation of the corresponding proteins. B) Significantly enriched Gene Ontology Biological Process and KEGG pathways revealed by Enrichr analysis of the protein expression up- (red shade) or down- (green shade) regulated in Optimal Transfectant condition compared Standard Transfectant condition at 72 hpt. In vertical axis is the terms, in horizontal axis is the transformed FDR (−log10PValue).

We conducted functional enrichment analysis on proteins exhibiting high and low abundance levels across all time points (24, 48, and 72 hpt), with fold changes of at least 1.5 and FDR-adjusted p-values less than 0.01. During the initial stages of AAV production at 24 hpt, various biological processes and KEGG pathways displayed distinctive regulation patterns.

Specifically, upregulated processes included Golgi vesicle-mediated transport (e.g., EXOC4, CCDC93, VTI1B), RNA metabolism (e.g., RNGTT, PCBP2, LAGE3), regulation of the cell cycle (e.g., HMGA2, TP53, UCHL5, SIRT2, MAP2K6), fatty acid biosynthesis (e.g., ERLIN1, SIRT2), and cellular stress responses (e.g., DNAJB6, NUP85, HSPA6).

Additionally, several other biosynthetic pathways were upregulated, such as oligosaccharide-lipid intermediate processes (e.g., ALG12, MPDU1), cellular lipid biosynthesis, and glycerol ether biosynthesis (e.g., FAR1, ERLIN1), at the 24 hpt time point (Supplementary dataset 2). In parallel, downregulated processes included DNA-templated transcription (e.g., MAZ, SSU72), autophagy (e.g., MTMR9, FOXK1, ATP6V1C1), cellular localization, and ubiquitin protein-dependent catabolic processes (e.g., UBE2C, RNF25, FBXO7).

Furthermore, KEGG database-defined pathways, like pyrimidine metabolism (e.g., TK1, DCK), homeostasis (e.g., FTH1, ATP6V1C1, FTL), membrane trafficking (e.g., KLC4, DCTN5, TBC1D13, CLTB, COPS7A, ARF5, KIF15), mitochondrial biogenesis (e.g., UBE2C, UBE2E3), and vesicle-mediated transport proteins related to interleukin signaling pathways (e.g., IL-2 and IL-5), were downregulated at the 24 hpt time point (Fig. 5, A-B & Supplementary dataset 2).

At the 48- (hpt) time point, we observed a substantial escalation in the expression of viral proteins and GFP compared to the previous 24 hpt time point. This heightened expression correlated with the upregulation of a diverse spectrum of biological processes and KEGG pathways, including cell division, DNA-templated transcription, homeostasis, viral RNA transcription, cytoskeleton organization, metabolic processes involving organonitrogen compounds, catabolic processes dependent on ubiquitin proteins, ribosomal biogenesis, cellular localization, and catabolic processes. Notably, proteins associated with processes like ubiquitin ligases, transferases, and ferritin complex proteins were found to be highly abundant at 48 hpt (Fig. 6, A-B).

This phase of the cell cycle aligns with the exponential mature phase of cell development, during which several physiological activities experienced downregulation. These activities encompassed the generation of metabolites, energy production, carboxylic acid biosynthesis, mitochondrial ATP synthesis coupled with electron transport, and cellular catabolic processes. Additionally, specific cellular protein complexes, including Mre11, BRCA1-C, oxidoreductases, and other mitochondrial protein complexes, exhibited a reduction in their expression levels.

At 72 hpt, with maximal viral protein and GFP production and expression, we observed alterations in molecular functions such as oxidoreductase, nucleosomal binding, and acyl-CoA dehydrogenase activities as they were upregulated. Biological processes related to mitochondrion organization, cellular lipid metabolic processes, organonitrogen compound biosynthetic processes, DNA packaging complexes, stress responses, and apoptosis pathways were all upregulated. At the late stage of the cell cycle (i.e., 72 hpt), cells reach maturity and undergo programmed cell death. Consequently, biological processes such as cell cycle/division, cellular regulations, translation, ribosomal biogenesis, chromosome segregation, and vesicle budding were all downregulated. Additionally, KEGG pathways such as Ubiquitin-mediated proteolysis and other signaling pathways experienced downregulation. Mitochondrial and membrane-bound proteins were also downregulated within cellular components at this stage of the cell cycle.

### Protein-Protein Interaction Network reveals role of HEK293T proteins in AAV Replication, Assembly, and Protein Synthesis

Our investigation shows the dynamic regulation of the host proteome, which plays a pivotal role in governing the processes of AAV2 replication, assembly, and protein expression. To systematically explore these proteomic changes, we conducted an exhaustive review of existing literature and performed comprehensive database searches, leading to the identification of around 124 host proteins exhibiting distinct expression patterns at various time points post-transfection (see Supplementary dataset 3).

To gain a more profound understanding of the functional implications of these proteins, we employed an extensive Protein-Protein Interaction Network Functional Enrichment Analysis using fullString network settings. Subsequently, we applied K-means clustering with a stringent confidence threshold of 0.700, which categorized these proteins into seven unique clusters (as illustrated in Fig. 8).

**Fig. 8.**
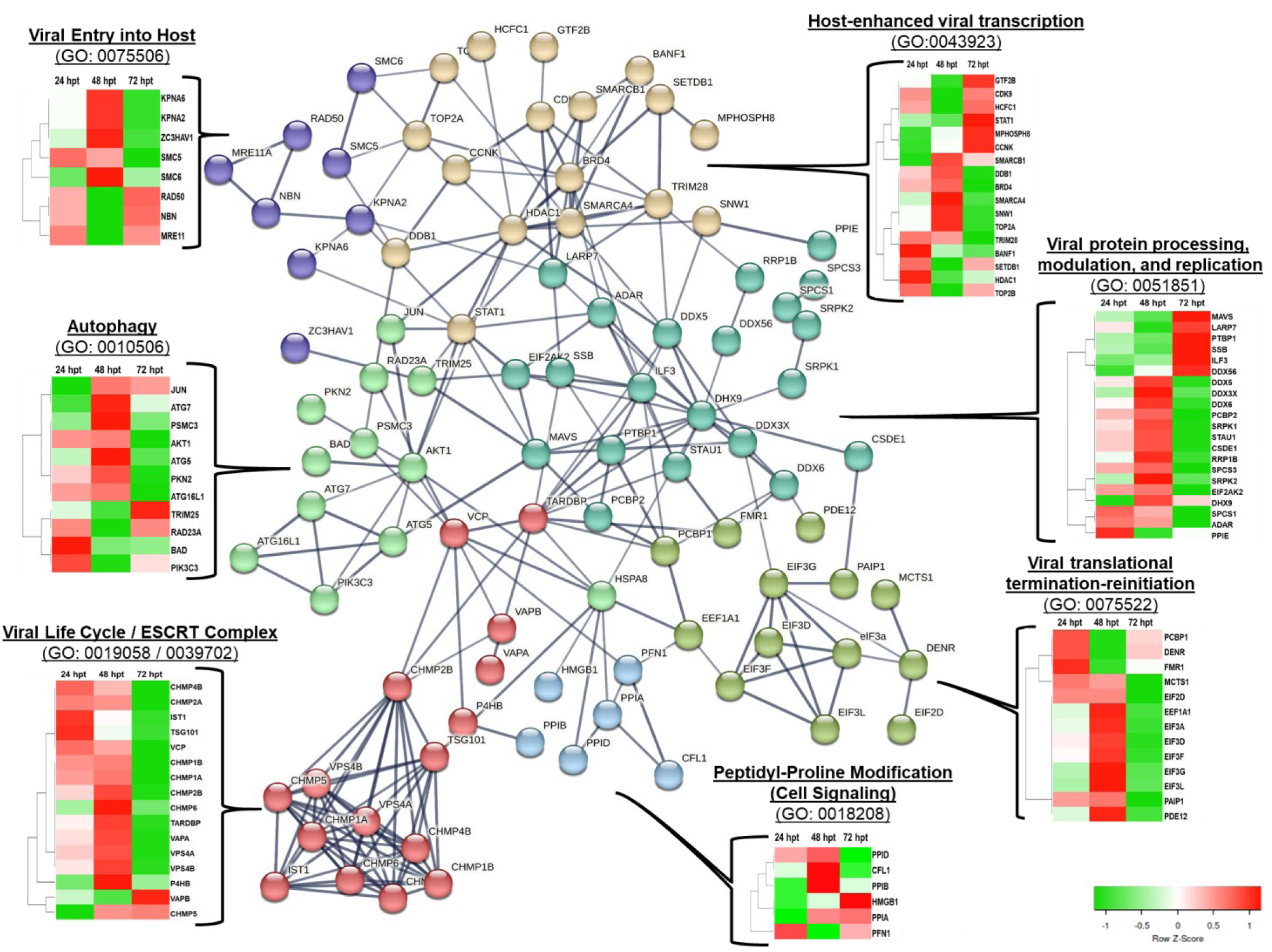
**Proteins Associated with AAV Replication and Assembly: Temporal Protein- Protein Interactions in HEK293T Cells.**

These seven clusters, comprised of 92 proteins (refer to Supplementary dataset 3 and Table 1, revealed associations with functional enrichment in specific pathways. These pathways encompassed Viral Entry into Host (GO: 0075506), Viral protein processing, modulation, and replication (GO: 0051851), Autophagy (GO: 0010506), Viral Life Cycle / ECRT Complex (GO: 0019058 / 0039702), Peptidyl-Proline Modification (Cell Signaling) (GO: 0018208), Viral translational termination-reinitiation (GO: 0075522), and Host-enhanced viral transcription (GO: 0043923). Notable examples include DNA repair proteins like MRE11 and RAD50, which played a role in negatively regulating viral entry into the host cell (GO: 0075506).

Throughout this study, we identified an enrichment in processes related to IRES-dependent viral translational initiation and the ESCRT complex, well-known for its involvement in cellular budding. This enrichment stemmed from the differential expression observed in a cluster of proteins, including Charged multivesicular body proteins (CHMP1A, CHMP1B, CHMP2A, CHMP2B, CHMP4B, CHMP5, CHMP6), Vesicle-associated membrane proteins (VAPA, VPS4A, VPS4B, VAPB), and other significant proteins like IST1, TSG101, VCP, TARDBP, and P4HB. The cluster associated with viral translational termination-reinitiation processes exhibited regulatory control by protein families, including Eukaryotic translation initiation factors, Poly(rC)-binding protein 1 (PCBP1), Density-regulated protein (DENR), Synaptic functional regulator (FMR1), Malignant T-cell-amplified sequence 1 (MCTS1), among others.

Our enrichment analysis further highlighted the differential expression of proteins involved in host cell signaling, encompassing Peptidyl-prolyl cis-trans isomerase D (PPIA, PPIB, PPID), Cofilin, non-muscle isoform (CFL1), High mobility group protein 1 (HMGB1), and Profilin- 1 (PFN1).

The observed variations in protein expression across these seven clusters collectively signify that, at 48 hpt, there is a notable increase in the abundance of host proteins that are typically found at lower levels during the early and late stages of cell growth (as depicted in Fig. 8).

## Discussion

This study was designed to provide an extensive exploration of the optimization of transfection conditions, with a focus on elevating the production of recombinant Adeno-Associated Virus (rAAV). Through the application of Design of Experiments (DoE), various critical factors influencing the transfection process, including cell density, DNA concentration, plasmid ratios, and PIE Max concentrations, were methodically fine-tuned to enhance efficiency. The implementation of three transfection conditions, denoted as Condition 1 (Standard), Condition 2 (Sub-optimal), and Condition 3 (Optimal), in conjunction with HEK293T cells cultured in FreeStyle™ F17 Expression Medium-293 media, was carried out with precision. The cells were seeded at a density of 1 × 10E6 viable cells/ml in 125 ml shaker flasks on the day of transfection.

A comprehensive proteome profiling approach was undertaken to gain insights into the molecular mechanisms associated with rAAV production. A spectral library was constructed using mass spectrometry data from HEK293T cells, both with and without AAV2 transfections, along with data from subcellular organelles. This library encompassed a diverse range of proteins, enabling the quantification of approximately 7,183 protein groups.

Subsequently, a comparative analysis was conducted across the three transfection conditions, revealing distinct proteome profiles at 24, 48, and 72 hours hpt. Principal Component Analysis (PCA) illustrated the temporal clustering of proteomes based on the infection time point. Differential expression analysis identified proteins with significant abundance changes, shedding light on the dynamic alterations in protein levels during AAV production.

Specifically, at 24 hpt, various biological processes and pathways associated with cellular processes, RNA metabolism, and cell cycle regulation exhibited upregulation. Conversely, pathways related to DNA-templated transcription, autophagy, and ubiquitin protein-dependent catabolic processes were downregulated. At 48 hpt, increased viral protein and GFP expression coincided with the upregulation of cell division, DNA-templated transcription, homeostasis, and various metabolic processes. Notably, proteins involved in ubiquitin ligases and ferritin complexes displayed high abundance. By 72 hpt, when viral protein expression peaked, upregulated functions included oxidoreductase activities, nucleosomal binding, and acyl-CoA dehydrogenase activities. Biological processes related to mitochondrion organization, cellular lipid metabolism, and apoptosis pathways were also upregulated, while cellular processes such as cell cycle/division, translation, and vesicle budding were downregulated. Further, the host cell’s innate immune response typically serves as the initial barrier against viral dissemination. It has been documented that, within the conventional transfection parameters applied to the HEK293 cell line, the host cell’s innate immune system tends to exhibit an upregulation^8,26–32^. However, in our case, when comparing our optimal transfection conditions to the standard ones, we observed a notable downregulation or reduced expression of proteins associated with immune response (such as ASAH1, CREB1, CD81, POLR3F, STOM, ATP11B, ATP6V1C1, NFKB2) across the time points (Supplementary Data GO Analysis). We also noticed higher expression of host cell proteins associated to import pathways during the peak rAAV production stages^33^. AAV encodes several proteins those are required for its replication. These replication proteins are involved in processes such as DNA replication and transcription. Importin proteins such as NUP107, NUP85, KPNA2, KPNA1 may facilitate the nuclear import of these viral replication proteins, allowing them to carry out their functions in the nucleus^34–36^ (Fig. 5-7 and Supplementary dataset 2).

It is noteworthy to emphasize that the AAV viral production cycle is critically dependent on the cellular replication and translation apparatus of the host cell. Given that viral proteins and viral nucleic acids are perceived as exogenous entities within the host cell milieu, their presence has the potential to trigger a series of host cell stress reactions, including but not limited to autophagy, endoplasmic reticulum (ER) stress, and programmed cell death (apoptosis). Our investigation unveiled an increased activation of these stress responses during phases associated with viral production^4,7,26,37–41^ (Fig. 5-7). Supplementary dataset 1 and 2 provides details of the GO enrichment results and the FC values for all the time-points post transfection for the Optimal transfection condition and list of host-cell proteins directly associated with viral replication and production.

In our ongoing investigation of the proteome, our focus shifted toward the identification of 124 host proteins exhibiting unique expression patterns during rAAV production. Through the utilization of Protein-Protein Interaction Network Functional Enrichment Analysis and K- means clustering, we effectively categorized these proteins into seven distinct clusters. Each cluster was closely linked with specific functional enrichment pathways, encompassing pivotal processes like Viral Entry into Host, Viral protein processing, modulation, and replication, Autophagy, Viral Life Cycle / ESCRT Complex, Peptidyl-Proline Modification, Viral translational termination-reinitiation, and Host-enhanced viral transcription.

It is worth noting that previous research has documented the multifaceted roles of the Endosomal Sorting Complex Required for Transport (ESCRT) machinery, which underpin various fundamental cellular processes, including cell cytokinesis, organelle and vesicle biogenesis, maintenance of nuclear-cytoplasmic compartmentalization, and endolysosomal activity^42–44^. In our specific study, we identified distinctive expression patterns of these proteins when compared to standard transfection conditions.

Furthermore, the core of the ESCRT machinery comprises three essential complexes, namely ESCRT-I, ESCRT-II, and ESCRT-III, along with the AAA-type ATP-ase, vacuolar protein sorting (VPS) 4^45–49^. Our investigation also uncovered variations in the expression of proteins within the CHMP family (Charged Multivesicular Body Proteins), including CHMP1B, CHMP1A, CHMP2A, CHMP2B, CHMP4B, CHMP5, and CHMP6^50,51^. These proteins play pivotal roles in processes such as endosomal sorting, multivesicular body (MVB) formation, and the budding of viruses, each contributing distinct functions within the ESCRT pathway^48,52–55^.

Additionally, we identified AAA+ ATPase and vacuolar protein sorting VPS4 proteins, specifically VPS4A and VPS4B, which have been previously recognized for their critical involvement in recycling ESCRT-III components and membrane remodelling^56,57^ (Refer to Supplementary dataset 3).

Interestingly, AAV also appears to exploit autophagy to aid in its escape from endosomes. The autophagy machinery, including components like autophagy-related proteins (ATGs) utilized by AAV to facilitate its transport from endosomes to the cytoplasm, where it can initiate productive infection. Apart from these RLR signaling pathway is the major sensing pathway for RNA viruses. Our study found that this pathway was enriched during the rAAV production (Supplementary dataset 3).

Certain peptidyl-proline modifications are known to play roles in cell signaling pathways, and we identified proteins like Peptidyl-prolyl cis-trans isomerases (PPIA, PPIB, PPID), Profilin-1 (PFN1), Cofilin-1 (CFL1), High mobility group protein B1 (HMGB1) with roles in regulating cellular responses.^58^.

In summary, this study offers comprehensive insights into the optimization of transfection conditions for enhanced rAAV production. By understanding the molecular functions of host cell proteins, we have shed light on the dynamic changes within the host proteome during AAV production. This not only provides a foundation for improving the efficiency of viral vector production but also highlights the potential for optimizing plasmid molar ratios and designing better HEK293T cell lines. With this knowledge, we can develop improved transient transfection conditions to achieve higher yields of AAV and other viral vectors for a wide range of applications.

## Materials and Methods

### Ethics Statement

All experiments were conducted in accordance with the approved guidelines of the Bioprocessing Technology Institute (BTI), Agency for Science, Technology and Research (A*STAR), Singapore.

### Experimental Design

In order to conduct functional and molecular characterization studies on various transient transfection conditions in the HEK293T cell line responsible for AAV2 production, we employed label-free quantitative proteomics using the SWATH-DIA method with a peptide- centric data analysis approach ^59^.Within this study, we investigated three distinct triple transfection conditions: the Standard condition (with a pHelper: pAAV_RC2: pAAV_GFP ratio of 1:1:1), a Sub-optimal condition, and the Optimal transfection condition. Sample collections were carried out at three different time points: 24 hours’ post-transfection (hpt), 48 hpt, and 72 hpt (as illustrated in Fig. 2). To ensure robust protein identification and consistent quantification using the SWATH technique, we meticulously constructed a comprehensive SWATH spectral library specific to the HEK293T cell line involved in AAV production (as depicted in Fig. 1). Subsequently, following deconvolution and spectral matching, we conducted a variety of statistical analyses.

### rAAV Production

In-house suspension-adapted HEK293T (ATCC) was transfected using pHelper, pAAV_RC2 and pAAV_GFP plasmids (CellBiolabs, VPK-402). Transfection was performed using PeiMax. Cells were cultured in F17 (ThermoFisher Scientific, A1383501) medium supplemented with 8 mM L-Glutamine (ThermoFisher Scientific, 25030081) and 0.1% Pluronic-F68 (ThermoFisher Scientific, 24040032). Cells were harvested for tittering 72 hours post-transfection. Design of Experiments (DoE) was used to optimize the rAAV transfection process.

### rAAV tittering

Cells were lysed and lysates were processed as described by Grieger, et al. ^19^. rAAV titering was performed on crude cell lysates using dye-based qPCR (NEB, M3003L) with primers against ITR regions. ITR primer sequences were obtained from Aurnhammer, et al. ^60^. Linearized plasmid standards were obtained by digesting pAAV_GFP with HindIII (NEB, R3104S) restriction enzyme.

### Sample Preparation

The cell pellet was lysed in lysis buffer (5% SDS, 50 mM TEAB, with benzonase) and clarified by centrifugation. Protein concentration was determined using the BCA Protein Assay kit.

100 µg of protein lysate was reduced and alkylated, trypsin-digested overnight at 37°C on an S-Trap mini spin column (ProtiFi) following manufacturer protocol. Eluted peptides were dried and reconstituted in loading buffer (1% formic acid and 2% acetonitrile) for mass spectrometry analysis.

### Data-dependent acquisition (DDA) – MS and HEK293T Spectral Library generation

For HEK293T spectral library construction, high-pH fractionated samples were analyzed using a TripleTOF 6600 (SCIEX) mass spectrometer in DDA mode, as previously reported ^61^. Raw DDA data files were processed with ProteinPilot^TM^ software (V5.0.2, SCIEX) against a human UniProt Proteome reference, with settings as previously described^61^. Result files from the search were used to generate input libraries for the spectral library in Spectronaut^TM^ software (Version: 17.6.230428.55965, Quasar) following recommended default settings. A secondary spectral library was created directly from DDA data files using the Pulsar database search engine in Spectronaut. Combining results from both ProteinPilot^TM^-based and Pulsar- based DDA searches enhanced library comprehensiveness, incorporating newly identified peptides into existing proteins and adding newly detected proteins to the base library.

### SWATH-DIA Data Acquisition

The SWATH-MS acquisition on TripleTOF 6600 (SCIEX) on DIA mode, the LC-MS instrument with specific settings as previously reported ^61^: a 100-variable window, a survey scan (MS1) from 350 to 1,250Lm/z for 50 ms, and high-sensitivity MS2 spectra from 100 to 1,500Lm/z for 25 ms. The total cycle time was approximately 2.5 s. Collision energy was applied to doubly charged precursors centered within the isolation window, using a collision energy equation like in DDA, with a collision energy spread (CES) of 5LeV.

### Statistical Analysis

Comparative statistical analysis of protein identification and quantification was performed for the transfection conditions, we used Spectronaut^TM^ software (Version: 17.0.221202.55965, Quasar) with our in-house comprehensive spectral library for HEK293T. The process of the functional classifications was conducted as follows. The first step was to cluster protein expression data using two software programs, Cluster 3.0 and Java Treeview for hireacrchical clustering and visualization of the results from proteome datasets. Clustered genes can later be decoded by Bulk Gene Searching Systems in Java (BGSSJ). BGSSJ is an XML-based Java application that systemizes lists of interesting genes and proteins for biological interpretation in the context of gene ontology3–5. Gene ontology analysis was performed using webgestalt, the filtering condition used was the Benjamini-Hochberg test with a p-value and FDR threshold of 0.05 performed using Over-Representation Analysis (ORA).

### Software

All analyses were performed in R (v.4.2.2) and open source tools, and the figures assembled in Adobe Illustrator. Fig. 1 and 2 was created with Biorender (cloud-based software).

## Reporting summary

Further information on research design is available in the Cell Portfolio Reporting Summary linked to this article.

## Data Availability

The personal data are not publicly available due to them containing information that could compromise research participant privacy. All other data are provided in the article and its Supplementary files or from the corresponding authors upon request. Source data are provided with this paper. All results described in the manuscript are presented in main or supplementary figures or datasets. All results from the statistical analysis are available as a shiny app resource at SRplot for data analysis and visualization. Data are available via ProteomeXchange with identifier PXD046574.

## Author Contributions

**ATP:** Conducted proteomics studies, overall data analysis, prepared figures and manuscript preparation, **ET**: Involved in design of experiment (DoE), conducted HEK293T cell culture, AAV titration experiments and reviewed the manuscript, **YJK**: prepared protein lysate and reviewed the manuscript, **SK** and **BX**: Supervised the study, review and mentorship. All authors have read and approved the final manuscript.

## References

1 Ail, D., Malki, H., Zin, E. A. & Dalkara, D. Adeno-Associated Virus (AAV) - Based Gene Therapies for Retinal Diseases: Where are We? Appl Clin Genet 16, 111–130, doi:10.2147/TACG.S383453 (2023).

2 Abulimiti, A., Lai, M. S. & Chang, R. C. Applications of adeno-associated virus vector- mediated gene delivery for neurodegenerative diseases and psychiatric diseases: Progress, advances, and challenges. Mech Ageing Dev 199, 111549, doi:10.1016/j.mad.2021.111549 (2021).

3 Brown, N., Song, L., Kollu, N. R. & Hirsch, M. L. Adeno-Associated Virus Vectors and Stem Cells: Friends or Foes? Hum Gene Ther 28, 450–463, doi:10.1089/hum.2017.038 (2017).

4 Aslanidi, G., Lamb, K. & Zolotukhin, S. An inducible system for highly efficient production of recombinant adeno-associated virus (rAAV) vectors in insect Sf9 cells. Proc Natl Acad Sci U S A 106, 5059–5064, doi:10.1073/pnas.0810614106 (2009).

5 Aucoin, M. G. et al. Virus-like particle and viral vector production using the baculovirus expression vector system/insect cell system: adeno-associated virus-based products. Methods Mol Biol 388, 281–296, doi:10.1007/978-1-59745-457-5_14 (2007).

6 Galibert, L. & Merten, O. W. Latest developments in the large-scale production of adeno- associated virus vectors in insect cells toward the treatment of neuromuscular diseases. J Invertebr Pathol 107 Suppl, S80-93, doi:10.1016/j.jip.2011.05.008 (2011).

7 Herrmann, A. K. et al. Impact of the Assembly-Activating Protein on Molecular Evolution of Synthetic Adeno-Associated Virus Capsids. Hum Gene Ther 30, 21–35, doi:10.1089/hum.2018.085 (2019).

8 Arjomandnejad, M., Dasgupta, I., Flotte, T. R. & Keeler, A. M. Immunogenicity of Recombinant Adeno-Associated Virus (AAV) Vectors for Gene Transfer. BioDrugs 37, 311–329, doi:10.1007/s40259-023-00585-7 (2023).

9 Bastola, P., Song, L., Gilger, B. C. & Hirsch, M. L. Adeno-Associated Virus Mediated Gene Therapy for Corneal Diseases. Pharmaceutics 12, doi:10.3390/pharmaceutics12080767 (2020).

10 Bera, A. & Sen, D. Promise of adeno-associated virus as a gene therapy vector for cardiovascular diseases. Heart Fail Rev 22, 795–823, doi:10.1007/s10741-017-9622-7 (2017).

11 Berns, K. I. & Srivastava, A. Next Generation of Adeno-Associated Virus Vectors for Gene Therapy for Human Liver Diseases. Gastroenterol Clin North Am 48, 319–330, doi:10.1016/j.gtc.2019.02.005 (2019).

12 Brommel, C. M., Cooney, A. L. & Sinn, P. L. Adeno-Associated Virus-Based Gene Therapy for Lifelong Correction of Genetic Disease. Hum Gene Ther 31, 985–995, doi:10.1089/hum.2020.138 (2020).

13 Buning, H., Perabo, L., Coutelle, O., Quadt-Humme, S. & Hallek, M. Recent developments in adeno-associated virus vector technology. J Gene Med 10, 717–733, doi:10.1002/jgm.1205 (2008).

14 Fu, Q., Polanco, A., Lee, Y. S. & Yoon, S. Critical challenges and advances in recombinant adeno-associated virus (rAAV) biomanufacturing. Biotechnol Bioeng 120, 2601–2621, doi:10.1002/bit.28412 (2023).

15 Joshi, P. R. H., Venereo-Sanchez, A., Chahal, P. S. & Kamen, A. A. Advancements in molecular design and bioprocessing of recombinant adeno-associated virus gene delivery vectors using the insect-cell baculovirus expression platform. Biotechnol J 16, e2000021, doi:10.1002/biot.202000021 (2021).

16 Ou, J. et al. Recent advances in upstream process development for production of recombinant adeno-associated virus. Biotechnol Bioeng, doi:10.1002/bit.28545 (2023).

17 Zhao, H. et al. Creation of a High-Yield AAV Vector Production Platform in Suspension Cells Using a Design-of-Experiment Approach. Mol Ther Methods Clin Dev 18, 312–320, doi:10.1016/j.omtm.2020.06.004 (2020).

18 Adamson-Small, L., Potter, M., Byrne, B. J. & Clement, N. Sodium Chloride Enhances Recombinant Adeno-Associated Virus Production in a Serum-Free Suspension Manufacturing Platform Using the Herpes Simplex Virus System. Hum Gene Ther Methods 28, 1–14, doi:10.1089/hgtb.2016.151 (2017).

19 Grieger, J. C., Soltys, S. M. & Samulski, R. J. Production of Recombinant Adeno-associated Virus Vectors Using Suspension HEK293 Cells and Continuous Harvest of Vector From the Culture Media for GMP FIX and FLT1 Clinical Vector. Mol Ther 24, 287–297, doi:10.1038/mt.2015.187 (2016).

20 Selvaraj, N. et al. Detailed Protocol for the Novel and Scalable Viral Vector Upstream Process for AAV Gene Therapy Manufacturing. Hum Gene Ther 32, 850–861, doi:10.1089/hum.2020.054 (2021).

21 Nguyen, T. N. T. et al. Mechanistic model for production of recombinant adeno-associated virus via triple transfection of HEK293 cells. Mol Ther Methods Clin Dev 21, 642–655, doi:10.1016/j.omtm.2021.04.006 (2021).

22 Li, J., Samulski, R. J. & Xiao, X. Role for highly regulated rep gene expression in adeno- associated virus vector production. J Virol 71, 5236–5243, doi:10.1128/JVI.71.7.5236-5243.1997 (1997).

23 Xiao, X., Li, J. & Samulski, R. J. Production of high-titer recombinant adeno-associated virus vectors in the absence of helper adenovirus. J Virol 72, 2224–2232, doi:10.1128/JVI.72.3.2224-2232.1998 (1998).

24 Strasser, L. et al. Proteomic Landscape of Adeno-Associated Virus (AAV)-Producing HEK293 Cells. Int J Mol Sci 22, doi:10.3390/ijms222111499 (2021).

25 Collins, B. C. et al. Multi-laboratory assessment of reproducibility, qualitative and quantitative performance of SWATH-mass spectrometry. Nat Commun 8, 291, doi:10.1038/s41467-017-00249-5 (2017).

26 Wang, Y., Fu, Q., Lee, Y. S., Sha, S. & Yoon, S. Transcriptomic features reveal molecular signatures associated with recombinant adeno-associated virus production in HEK293 cells. Biotechnol Prog 39, e3346, doi:10.1002/btpr.3346 (2023).

27 Balakrishnan, B. et al. Activation of the cellular unfolded protein response by recombinant adeno-associated virus vectors. PLoS One 8, e53845, doi:10.1371/journal.pone.0053845 (2013).

28 Calcedo, R., Chichester, J. A. & Wilson, J. M. Assessment of Humoral, Innate, and T-Cell Immune Responses to Adeno-Associated Virus Vectors. Hum Gene Ther Methods 29, 86–95, doi:10.1089/hgtb.2018.038 (2018).

29 Coroadinha, A. S. Host Cell Restriction Factors Blocking Efficient Vector Transduction: Challenges in Lentiviral and Adeno-Associated Vector Based Gene Therapies. Cells 12, doi:10.3390/cells12050732 (2023).

30 Dauletbekov, D. L., Pfromm, J. K., Fritz, A. K. & Fischer, M. D. Innate Immune Response Following AAV Administration. Adv Exp Med Biol 1185, 165–168, doi:10.1007/978-3-030-27378-1_27 (2019).

31 Hamilton, B. A. & Wright, J. F. Challenges Posed by Immune Responses to AAV Vectors: Addressing Root Causes. Front Immunol 12, 675897, doi:10.3389/fimmu.2021.675897 (2021).

32 Hareendran, S. et al. Adeno-associated virus (AAV) vectors in gene therapy: immune challenges and strategies to circumvent them. Rev Med Virol 23, 399–413, doi:10.1002/rmv.1762 (2013).

33 Nicolson, S. C. & Samulski, R. J. Recombinant adeno-associated virus utilizes host cell nuclear import machinery to enter the nucleus. J Virol 88, 4132–4144, doi:10.1128/JVI.02660-13 (2014).

34 Weinberg, M. S. et al. Recombinant adeno-associated virus utilizes cell-specific infectious entry mechanisms. J Virol 88, 12472–12484, doi:10.1128/JVI.01971-14 (2014).

35 Whittaker, G. R. & Helenius, A. Nuclear import and export of viruses and virus genomes. Virology 246, 1–23, doi:10.1006/viro.1998.9165 (1998).

36 Whittaker, G. R., Kann, M. & Helenius, A. Viral entry into the nucleus. Annu Rev Cell Dev Biol 16, 627–651, doi:10.1146/annurev.cellbio.16.1.627 (2000).

37 Aponte-Ubillus, J. J. et al. Proteome profiling and vector yield optimization in a recombinant adeno-associated virus-producing yeast model. Microbiologyopen 9, e1136, doi:10.1002/mbo3.1136 (2020).

38 Destro, F. et al. Mechanistic modeling explains the production dynamics of recombinant adeno-associated virus with the baculovirus expression vector system. Mol Ther Methods Clin Dev 30, 122–146, doi:10.1016/j.omtm.2023.05.019 (2023).

39 Gallo-Ramirez, L. E., Ramirez, O. T. & Palomares, L. A. Intracellular localization of adeno- associated viral proteins expressed in insect cells. Biotechnol Prog 27, 483–493, doi:10.1002/btpr.565 (2011).

40 Li, L. et al. Production and characterization of novel recombinant adeno-associated virus replicative-form genomes: a eukaryotic source of DNA for gene transfer. PLoS One 8, e69879, doi:10.1371/journal.pone.0069879 (2013).

41 Ruffing, M., Zentgraf, H. & Kleinschmidt, J. A. Assembly of viruslike particles by recombinant structural proteins of adeno-associated virus type 2 in insect cells. J Virol 66, 6922–6930, doi:10.1128/JVI.66.12.6922-6930.1992 (1992).

42 Calistri, A., Reale, A., Palù, G. & Parolin, C. Why Cells and Viruses Cannot Survive without an ESCRT. 10, 483 (2021).

43 Miller, S. & Krijnse-Locker, J. Modification of intracellular membrane structures for virus replication. Nat Rev Microbiol 6, 363–374, doi:10.1038/nrmicro1890 (2008).

44 Spuul, P. et al. Assembly of alphavirus replication complexes from RNA and protein components in a novel trans-replication system in mammalian cells. J Virol 85, 4739–4751, doi:10.1128/JVI.00085-11 (2011).

45 Barajas, D., Kovalev, N., Qin, J. & Nagy, P. D. Novel mechanism of regulation of tomato bushy stunt virus replication by cellular WW-domain proteins. J Virol 89, 2064–2079, doi:10.1128/JVI.02719-14 (2015).

46 Campsteijn, C., Vietri, M. & Stenmark, H. Novel ESCRT functions in cell biology: spiraling out of control? Curr Opin Cell Biol 41, 1–8, doi:10.1016/j.ceb.2016.03.008 (2016).

47 Hurley, J. H. ESCRTs are everywhere. EMBO J 34, 2398–2407, doi:10.15252/embj.201592484 (2015).

48 Jiang, B., Himmelsbach, K., Ren, H., Boller, K. & Hildt, E. Subviral Hepatitis B Virus Filaments, like Infectious Viral Particles, Are Released via Multivesicular Bodies. J Virol 90, 3330–3341, doi:10.1128/JVI.03109-15 (2015).

49 Shields, S. B. & Piper, R. C. How ubiquitin functions with ESCRTs. Traffic 12, 1306–1317, doi:10.1111/j.1600-0854.2011.01242.x (2011).

50 Robinson, M., Schor, S., Barouch-Bentov, R. & Einav, S. Viral journeys on the intracellular highways. Cell Mol Life Sci 75, 3693–3714, doi:10.1007/s00018-018-2882-0 (2018).

51 Isono, E. ESCRT Is a Great Sealer: Non-Endosomal Function of the ESCRT Machinery in Membrane Repair and Autophagy. Plant Cell Physiol 62, 766–774, doi:10.1093/pcp/pcab045 (2021).

52 Agromayor, M. et al. Essential role of hIST1 in cytokinesis. Mol Biol Cell 20, 1374–1387, doi:10.1091/mbc.e08-05-0474 (2009).

53 Ariumi, Y. et al. The ESCRT system is required for hepatitis C virus production. PLoS One 6, e14517, doi:10.1371/journal.pone.0014517 (2011).

54 Chen, B. J. & Lamb, R. A. Mechanisms for enveloped virus budding: can some viruses do without an ESCRT? Virology 372, 221–232, doi:10.1016/j.virol.2007.11.008 (2008).

55 Wegner, C. S., Rodahl, L. M. & Stenmark, H. ESCRT proteins and cell signalling. Traffic 12, 1291–1297, doi:10.1111/j.1600-0854.2011.01210.x (2011).

56 Corless, L., Crump, C. M., Griffin, S. D. & Harris, M. Vps4 and the ESCRT-III complex are required for the release of infectious hepatitis C virus particles. J Gen Virol 91, 362–372, doi:10.1099/vir.0.017285-0 (2010).

57 Han, H. & Hill, C. P. Structure and mechanism of the ESCRT pathway AAA+ ATPase Vps4. Biochem Soc Trans 47, 37–45, doi:10.1042/BST20180260 (2019).

58 Ren, X. & Hurley, J. H. Proline-rich regions and motifs in trafficking: from ESCRT interaction to viral exploitation. Traffic 12, 1282–1290, doi:10.1111/j.1600-0854.2011.01208.x (2011).

59 Schubert, O. T. et al. Building high-quality assay libraries for targeted analysis of SWATH MS data. Nat Protoc 10, 426–441, doi:10.1038/nprot.2015.015 (2015).

60 Aurnhammer, C. et al. Universal real-time PCR for the detection and quantification of adeno- associated virus serotype 2-derived inverted terminal repeat sequences. Hum Gene Ther Methods 23, 18–28, doi:10.1089/hgtb.2011.034 (2012).

61 Sim, K. H. et al. A comprehensive CHO SWATH-MS spectral library for robust quantitative profiling of 10,000 proteins. Sci Data 7, 263, doi:10.1038/s41597-020-00594-z (2020).

