## Supplementary material for "Temporal Insights into Molecular and Cellular Responses during rAAV Production in HEK293T Cells": Table 1

**Table 1:** Functionally Enriched Pathways and HEK293T Host Cell Genes/Proteins Implicated in Viral Replication, Assembly, and Protein Expression.

| **Functional Enrichment** | **Gene ID** | **Name** |
| --- | --- | --- |
| **Autophagy**  **(GO: 0010506)** | JUN | Transcription factor AP-1 |
|  | ATG7 | Ubiquitin-like modifier-activating enzyme ATG7 |
|  | PSMC3 | 26S proteasome regulatory subunit 6A |
|  | AKT1 | RAC-alpha serine/threonine-protein kinase |
|  | ATG5 | Autophagy protein 5 |
|  | PKN2 | Serine/threonine-protein kinase N2 |
|  | ATG16L1 | ATG16 autophagy related 16-like 1 |
|  | TRIM25 | E3 ubiquitin/ISG15 ligase TRIM25 |
|  | RAD23A | UV excision repair protein RAD23 |
|  | BAD | BCL2-antagonist of cell death, isoform CRA_c |
|  | PIK3C3 | Phosphatidylinositol 3-kinase catalytic subunit type 3 |
| **Viral Entry into Host**  **(GO: 0075506)** | KPNA6 | Importin subunit alpha-7 |
|  | KPNA2 | Importin subunit alpha-1 |
|  | ZC3HAV1 | Zinc finger CCCH-type antiviral protein 1 |
|  | SMC5 | Structural maintenance of chromosomes protein 5 |
|  | SMC6 | Structural maintenance of chromosomes 6 |
|  | RAD50 | DNA repair protein RAD50 |
|  | NBN | Nibrin |
|  | MRE11 | MRE11 homolog, double strand break repair nuclease |
| **Viral protein processing, modulation, and replication**  **(GO: 0051851)** | MAVS | Mitochondrial antiviral-signaling protein |
|  | LARP7 | La-related protein 7 |
|  | PTBP1 | Polypyrimidine tract-binding protein 1 |
|  | SSB | Autoantigen La (Fragment) |
|  | ILF3 | Interleukin enhancer binding factor 3, 90kDa, isoform CRA_d |
|  | DDX56 | RNA helicase (Fragment) |
|  | DDX5 | DEAD box protein 5 (Fragment) |
|  | DDX3X | RNA helicase |
|  | DDX6 | Probable ATP-dependent RNA helicase DDX6 |
|  | PCBP2 | Poly(rC)-binding protein 2 |
|  | SRPK1 | SRSF protein kinase 1 (Fragment) |
|  | STAU1 | Double-stranded RNA-binding protein Staufen homolog 1 |
|  | CSDE1 | Cold shock domain containing E1, RNA-binding |
|  | RRP1B | Ribosomal RNA processing protein 1 homolog B |
|  | SPCS3 | Signal peptidase complex subunit 3 |
|  | SRPK2 | SFRS protein kinase 2 |
|  | EIF2AK2 | Interferon-induced, double-stranded RNA-activated protein kinase |
|  | DHX9 | ATP-dependent RNA helicase A |
|  | SPCS1 | Signal peptidase complex subunit 1 |
|  | ADAR | Double-stranded RNA-specific adenosine deaminase |
|  | PPIE | Peptidyl-prolyl cis-trans isomerase E |
| **Viral Life Cycle / ECRT Complex (GO : 0019058 / 0039702)** | CHMP4B | Charged multivesicular body protein 4b |
|  | CHMP2A | Chromatin modifying protein 2A, isoform CRA_b |
|  | IST1 | IST1 homolog |
|  | TSG101 | Tumor susceptibility gene 101 protein |
|  | VCP | Transitional endoplasmic reticulum ATPase |
|  | CHMP1B | Charged multivesicular body protein 1b |
|  | CHMP1A | Charged multivesicular body protein 1a |
|  | CHMP2B | Charged multivesicular body protein 2b |
|  | CHMP6 | Charged multivesicular body protein 6 (Fragment) |
|  | TARDBP | TAR DNA-binding protein 43 |
|  | VAPA | Vesicle-associated membrane protein-associated protein A |
|  | VPS4A | Vesicle-fusing ATPase 4A |
|  | VPS4B | Vesicle-fusing ATPase 4B |
|  | P4HB | Protein disulfide-isomerase |
|  | VAPB | Vesicle-associated membrane protein-associated protein B/C |
|  | CHMP5 | Charged multivesicular body protein 5 |
| **Peptidyl-Proline Modification (Cell Signaling)**  **(GO: 0018208)** | PPID | Peptidyl-prolyl cis-trans isomerase D |
|  | CFL1 | Cofilin, non-muscle isoform |
|  | PPIB | Peptidyl-prolyl cis-trans isomerase B |
|  | HMGB1 | High mobility group protein 1 |
|  | PPIA | Peptidyl-prolyl cis-trans isomerase |
|  | PFN1 | Profilin-1 |
| **Viral translational termination-reinitiation**  **(GO: 0075522)** | PCBP1 | Poly(rC)-binding protein 1 |
|  | DENR | Density-regulated protein |
|  | FMR1 | Synaptic functional regulator FMR1 |
|  | MCTS1 | Malignant T-cell-amplified sequence 1 |
|  | EIF2D | Eukaryotic translation initiation factor 2D |
|  | EEF1A1 | Elongation factor 1-alpha 1 |
|  | EIF3A | Eukaryotic translation initiation factor 3 subunit A |
|  | EIF3D | Eukaryotic translation initiation factor 3 subunit D |
|  | EIF3F | Eukaryotic translation initiation factor 3 subunit F |
|  | EIF3G | Eukaryotic translation initiation factor 3 subunit G |
|  | EIF3L | Eukaryotic translation initiation factor 3 subunit L |
|  | PAIP1 | Polyadenylate-binding protein-interacting protein 1 |
|  | PDE12 | 2',5'-phosphodiesterase 12 |
| **Host-enhanced viral transcription**  **(GO: 0043923)** | GTF2B | Transcription initiation factor IIB |
|  | CDK9 | Cyclin-dependent kinase 9 |
|  | HCFC1 | Host cell factor 1 |
|  | STAT1 | Signal transducer and activator of transcription |
|  | MPHOSPH8 | M-phase phosphoprotein 8 |
|  | CCNK | Cyclin K, isoform CRA_c |
|  | SMARCB1 | SWI/SNF-related matrix-associated actin-dependent regulator of chromatin subfamily B member 1 |
|  | DDB1 | DNA damage-binding protein 1 |
|  | BRD4 | Bromodomain-containing protein 4 |
|  | SMARCA4 | Transcription activator BRG1 |
|  | SNW1 | SNW domain-containing protein 1 |
|  | TOP2A | DNA topoisomerase 2-alpha |
|  | TRIM28 | Transcription intermediary factor 1-beta |
|  | BANF1 | Barrier to autointegration factor 1, isoform CRA_a |
|  | SETDB1 | Histone-lysine N-methyltransferase SETDB1 |
|  | HDAC1 | Histone deacetylase 1 |
|  | TOP2B | DNA topoisomerase 2-beta |
